# Individual and social contributions to learning of mice in Intellicages: the general framework and mean field theory

**DOI:** 10.1101/2025.11.12.687796

**Authors:** Bartosz Jura, Michał Lenarczyk, Zofia Harda, Łukasz Szumiec, Magdalena Ziemiańska, Róż Kłosowicz, Jan Rodriguez Parkitna, Daniel Krzysztof Wójcik

## Abstract

Reinforcement learning enables the adaptation of behaviors to changes in the environment. Models of reinforcement learning based on using previous experience to guide future choices enable balancing exploitation of opportunities and exploring the options and provide accurate predictions of animal behavior in simple instrumental tasks. However, they fail to account for the critical advantage we and other animals have, namely the ability not only to learn from our own choices but also from observing the outcomes of choices made by others. Here, we propose a new conceptual, analytical, and computational framework combining point processes with reinforcement learning models for the description, analysis and modeling of individual and group effects of mice learning reward placement. We show that marked point processes provide a natural language to describe behavior of group housed mice. We show how different reinforcement models of the behavior, including group effects, can be studied effectively in this framework, using example experiments. With this framework we show that the group effects (peer pressure) are twice stronger than individual learning.

## Introduction

Learning from experience is a fundamental property of adaptive behavior, enabling organisms to modify future actions based on past outcomes. Reinforcement learning (RL) provides a computational framework for understanding this process by formalizing how actions are selected, evaluated, and updated in response to rewards and punishments^1^. At the neural level, RL theories have been tightly linked to the dopaminergic system, where phasic dopamine activity encodes prediction errors—the discrepancy between expected and received outcomes—that drives learning ^2–4^. Such mechanisms explain how value representations are dynamically updated across cortical and subcortical regions to guide choice behavior ^5;6^. Experiments show that human and animal decision-making strategies align well with temporal-difference algorithms^7;8^. These results have not only advanced our understanding of learning and decision-making but also implicated the relevance of RL models to neuropsychiatric conditions characterized by maladaptive choice behavior ^9;10^.

In many species, animals are able to learn not only from own experiences but also by observing and interacting with peers, which we call social learning ^11;12^. Until recently, most research on animal learning ignored the social aspects to avoid the complexity of several interacting individuals. If social behavior was studied, this usually involved either detailed observation of animal pairs or collective behavior of large groups ^13–15^. Recently, the need to study intermediate groups, where the effects of individuality and group interactions can be comparable, has been indicated ^16–22^. Another motivation for studies of medium sized cohorts came from the need to increase throughput of phenotypization. Genetically modified mice carrying mutations of genes expressed in the nervous system facilitate study of the relations between the gene and behavior. These relations are assessed with dedicated behavioral tests ^23^. To increase throughput of phenotypization several automated cage types were developed to allow simultaneous studies of mouse cohorts of different sizes^15–20;24^. This approach provided the group context for the behavioral studies, but also increased throughput and decreased the need for human handling, thus bringing more ecological conditions to the study. However, a systematic quantitative study of how animals learn in cohorts taking into account the aspects of individual and social learning is still missing.

Here we propose a theoretical and computational framework for modeling behavior of animal cohorts where the animal tracking is based on discrete events in time ^16–20;24^, which allows testing hypotheses on different individual and social learning strategies^25;26^. We illustrate this framework with data coming from the Intellicage system ^16;17;23;27^ where two groups, a majority and minority, learn reward placement available at different corners. The challenge for the animals is that both the reward placement and the group assignment change in time. If the animals learned only from their own experience the group assignment would be irrelevant. On the other hand, if the animals followed the prevailing choices in the group ^12;25^, those in the minority group (2 animals) should be more influenced by the behavior of the majority (12 animals). Using our framework we show that a model combining individual learning with following the group describes our data well and indicates most likely learning strategies used by individual animals, but also allows to quantify the relative importance of acting on one’s own experience as compared to following the group.

## Results

### Overview of the proposed framework

Our main result is a conceptual and computational framework for the description of behavior of interacting animal cohorts which are monitored by discrete events. We illustrate this framework with data from an experiment on mice learning reward placement in an Intellicage, hence we place our presentation in this context. Here we present an overview of our framework leaving the details to the Methods section.

The Intellicage is a rectangular cage with four automated learning chambers (corners), Fig. 1a. Every corner has two bottles which can be accessed by a mouse one at a time. The bottles can be used to provide the animal with water, rewarding saccharine solution, aversive quinine solution, addictive alcohol, etc., allowing for a multitude of experimental protocols to be scaled up from individual animal studies to long-term, more ecological context^27^. Up to 16 mice can be co-housed in an Intellicage ^23^. Every mouse has an RFID transponder placed subcutaneously which allows the mouse to be identified when interacting with the corners. This identification is used to annotate the recorded results of the experiment but also to design experiments where access to selected bottles may be allowed to specific animals.

**Figure 1:**
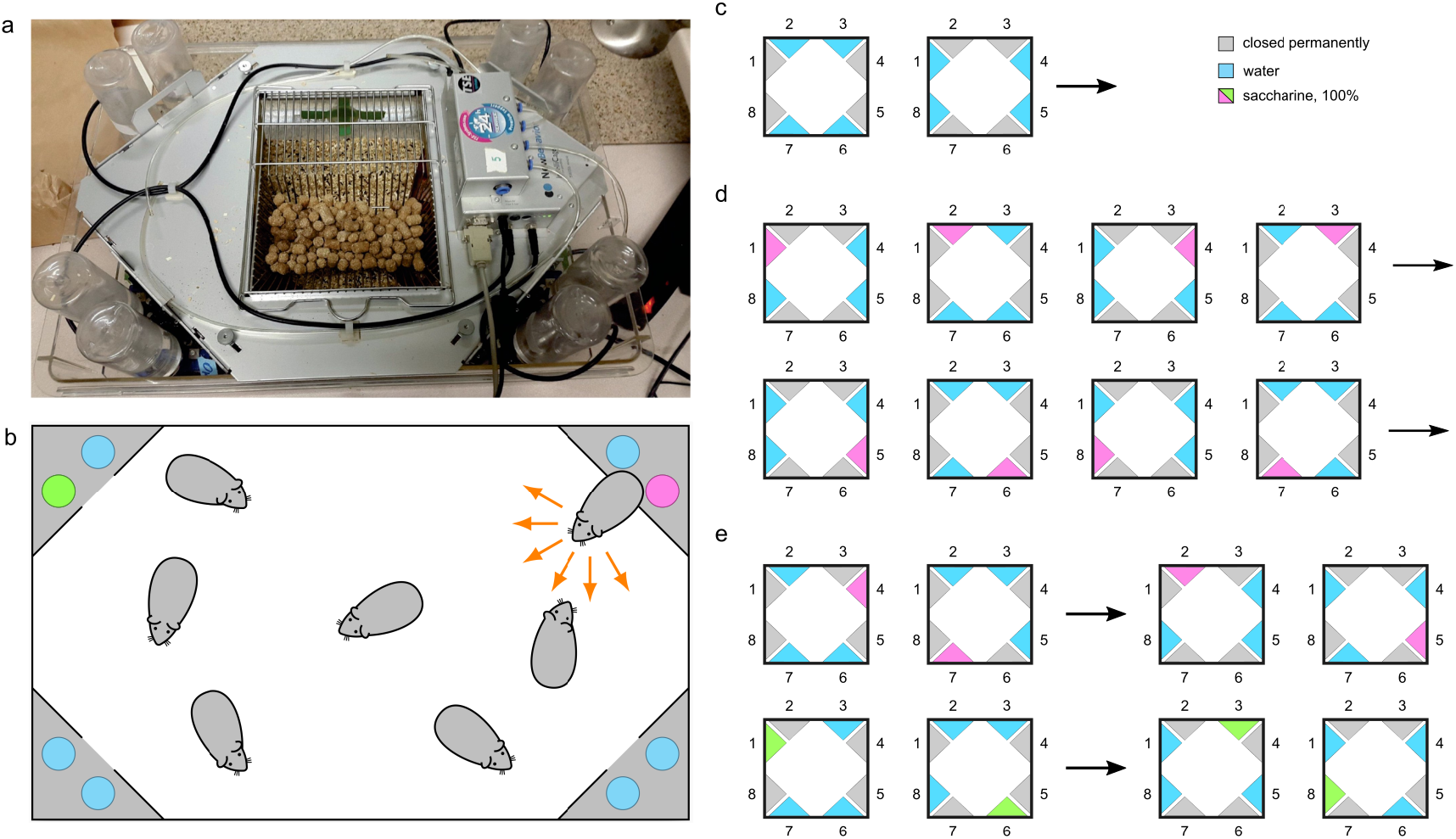
Intellicage design and experimental setup. **a** A photo and **b** a schema of the Intellicage System. **c–e** Experimental design. **c** 4 days of adaptation to the cage were followed by **d** 8 days with reward delivered to all the mice through the same bottle with the bottle location changing every 24 hours. **e** Experimental phase consisted of 48h periods in which a minority group consisting of 2 mice was offered access to saccharine in one corner while the rest (majority group) in another. Every 48h rewarded corners changed. Every 96h assignment to the minority group changed. Overall, every mouse was exactly once in the minority.

To describe the behavior of a cohort of mice in an Intellicage note that at any given time a mouse can be in one of the corners numbered 1 to 4 or in the central area which we call the “arena”. We can thus attribute to every mouse at each time a “mark” or “state” 0 to 4 that represents the space the mouse occupies. A mouse changes its state at discrete moments, therefore, a complete description of the mouse behavior that we can form based on available sensors, or its trajectory, is a collection of times when the mouse enters or leaves specific corners with corresponding corner numbers. Such an object is formally a path in a marked point process ^28^.

We assume that the mice explore the cage and visit corners randomly but the chances to visit each corner can be different and change in time. As a mouse explores the cage it may receive rewards or punishments in different corners which may affect how it values the corners and modify the probability of the visit. Also, the mice are aware of one another and peer pressure or group behavior may affect the choices made.

Identifying a true model of mouse decision making, taking into account learning by direct experience within corners but also indirect effects of peer behavior on the choices, is difficult. Instead we propose several probabilistic models, with some taking into account social transmission of knowledge, and compare their likelihoods given available data.

We assume every mouse keeps its own valuation of every corner which it builds during corner visits. The simplest learning model we consider is RW-learning^29^, where a mouse keeps a value *Q*_*a*_(*n*) of every corner *a* which is updated after every visit according to Rescorla-Wagner rule: *Q*_*a*_(*n* + 1) = *Q*_*a*_(*n*) + *α*(*R*_*n*_ − *Q*_*a*_(*n*)), where *α* is learning rate, *R*_*n*_ is the reward received on *n*-th visit, which is 0 for water, 1 for saccharine solution.

We also assume the mice keep track of the behavior of their mates and build a proxy model for socially attractive corners as experimental proportions of corner visits in the past. We assume every mouse makes a choice with some fixed probability based on what it learned, otherwise it uses the social model. The length of the observation window and the probability of following the crowd are parameters to be fit from the data.

The general probabilistic model to describe mice behavior in an Intellicage which we propose here is given by the probability *P* of transition of mouse *i* from area *j* to area *k* during time [*t, t* + *dt*] given the history of all mice in this cage up to the present time, *H*(*t*), or more precisely, by intensity of transitions, *λ*:

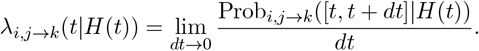

This can be decomposed as

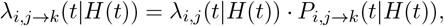

where *λ*_*i,j*_(*t*|*H*(*t*)) is the hazard function generating a transition of mouse *i* which at time *t* is in state *j*, and *P*_*i,j*→*k*_(*t*|*H*(*t*)) is the conditional probability of a transition from *j* to *k* at time *t* given complete history of all mice behavior.

The transition probability *P*_*i*,0→*k*_ depends on the self-learned value of individual corners and on the group behavior. The self-learned part is modeled by soft-max function of the valuation function *Q*_*i,k*_(*t*):

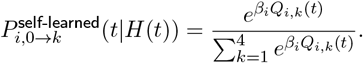

We considered five different models of self-learning which we call Base, Dual Alpha, Fictitious Update, Hybrid, and Four Alpha. Each of them realizes a learning process which leads to evolution of corner valuations in response to rewards provided during cage exploration.

We further assume that with probability 1 − *γ* the mouse makes a decision based on its own experience, and with probability *γ* the mouse selects a corner depending on the history of visits of all the mice. Hence, for *γ* = 0 we have models of self-learning where the social influence is neglected, and for *γ* = 1 we have models where a mouse ignores rewards and follows the crowd. Combining self-learning and group learning we obtain our model probability of a jump as

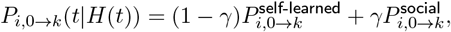

where 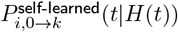 is the soft-max for one of the above models, and 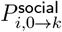 represents the mean behavior in the group in recent past characterized by the time-scale of social observation, *τ*_*s*_.

Under these assumptions the log-likelihood of observing a mouse trajectory splits into a sum of terms dependent only on the hazard function and terms dependent only on the learning model, which can be optimized independently. In the present work we focus on the second part which describes learning.

### Social learning dominates self-learning

To illustrate the power and practicality of the proposed framework we used it to disentangle individual and social effects in wild type mice C57BL/6 learning reward placement in Intellicages, and to understand their relative importance, in an experiment on a cohort of 14 female mice, Fig. 1. After adaptation the mice were divided into “majority” and “minority” groups consisting of 12 and 2 individuals, respectively. In every corner each animal had access to exactly one bottle, a saccharin solution in one corner, water in the rest. The rewarded bottles were located in different corners for different groups. Thus, if social cues played a role in learning the placement of the saccharin bottle, then every animal would be cued by others from its group and misdirected with regard to the position of the reward by the behavior of the other group. This misleading effect should be stronger for the minority group. The assignment of animals to groups changed every 4 days, in such a way that eventually every animal was a member of a minority group for exactly one 4-day period.

To facilitate understanding of our formal analysis which follows we first visually explore the data from the first half of the first experimental phase (E1), Fig. 2. Here, the first two mice (1 and 2) are in the minority group while the rest are in the majority group. Fig. 2**a, b** show animal activity as raster plots for the majority and minority groups, respectively. Each row represents a mouse, each point represents the animal entry to the corner, color codes the corner number. Red denotes the rewarded corner for the majority group (here: corner 1); blue denotes the rewarded corner for the minority group (here: corner 2). In this case, all animals learn the location of the reward, although the performance of animal 13 indicates it was not ∗ able to learn the position of the bottles with saccharin solution (marked by in Fig. 2**a**). Fig. 2**c, d** show average daily animal activity in groups while Fig. 2**e, f** show conditional probabilities to enter a given corner for a given population with color-coded corners. Probabilities and conditional probabilities were computed in moving rectangular windows of length 6h. Here red represents corner 1, which was rewarded for the majority, and blue represents corner 2, which was rewarded for the minority. We see that both groups learned the locations of the respective rewarded corners which is indicated by higher probability of making visits there and is marked with filled dots in Fig. 2**e, f**. Direct comparison of the group performance, however, shows that 1) the probability to go to the rewarded corner for a mouse in the majority group is somewhat higher than for the minority group (Fig. 2**g**); 2) the chances to go to the other group rewarded corner are higher for the minority group (Fig. 2**h**, red open circles), than for the majority group (Fig. 2**h**, blue open circles). This is consistent with the conjecture that the behavior of majority affects the behavior of minority stronger than the opposite.

**Figure 2:**
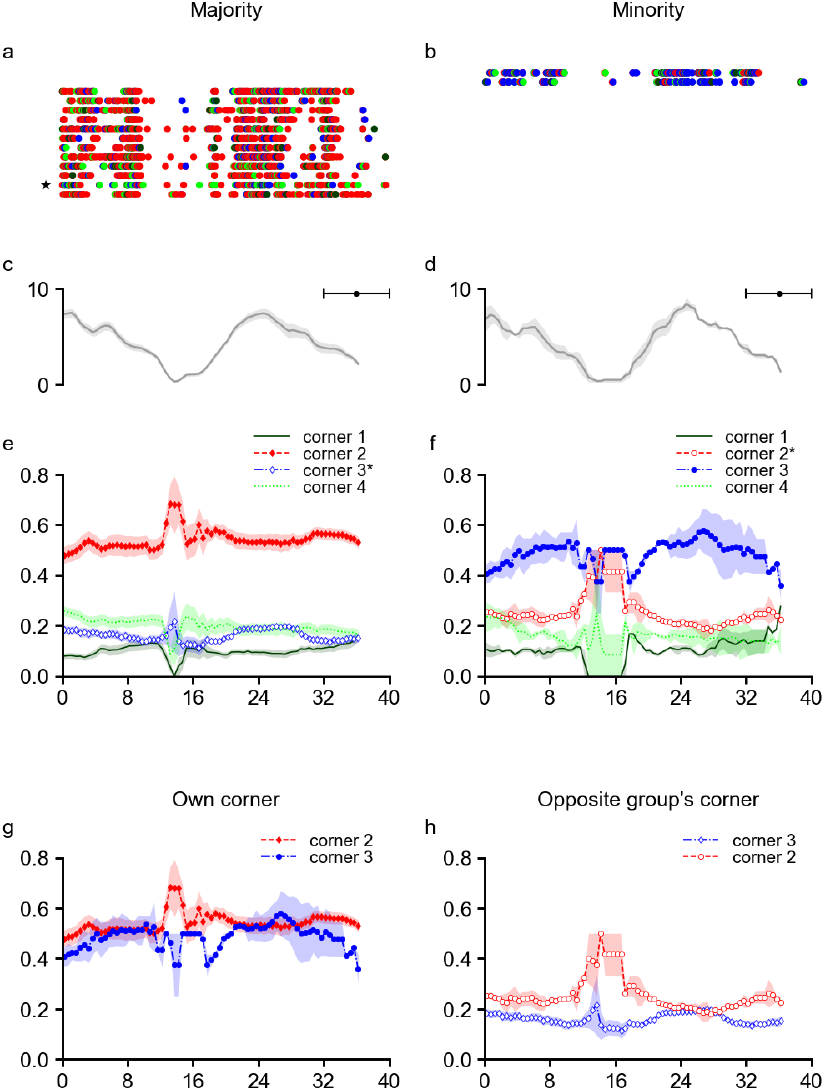
Representative results from the first half of the first experimental phase (E1). Here, the first two mice (1 and 2) are in the minority group while the rest are in the majority group. **a**,**b** animal activity as raster plots. Each point represents animal entry to the corner, color codes corner number. Red denotes the rewarded corner for the majority group (here: corner 1); blue denotes rewarded corner for the minority group (here: corner 2). **c, d** daily activity of groups. **e, f** conditional probabilities to enter a given corner for a given population. Filled dots represent the visits to group’s own reward corner, open dots represent the visits to the other group’s reward corner. **g** visits to both groups’ own reward corners. **h** visits of both groups to corners with reward for the other group.

Fig. 3 shows complete behavioral data from all the experimental phases for an example, mouse 14 (Fig. 3**a**–**c**, which was representative for the studied cohort. Breaks in the data represent malfunction of the recording apparatus and do not significantly affect the analysis. Fig. 3**a** shows the raster plot of mouse activity. Fig. 3**b** shows the valuations learned by the mouse. Fig. 3**c** shows the soft-max probabilities to enter the corner given the valuation. Note that only the self-learning part is shown here. Notice the lower probability to visit the rewarded corner of in the last experimental phase. The reason is that mouse 14 is in the minority group, hence group cues are misleading. But even if mouse 14 was able to identify its minority peer (mouse 13) this would not be helpful as mouse 13 is the worst performer of all (not shown).

**Figure 3:**
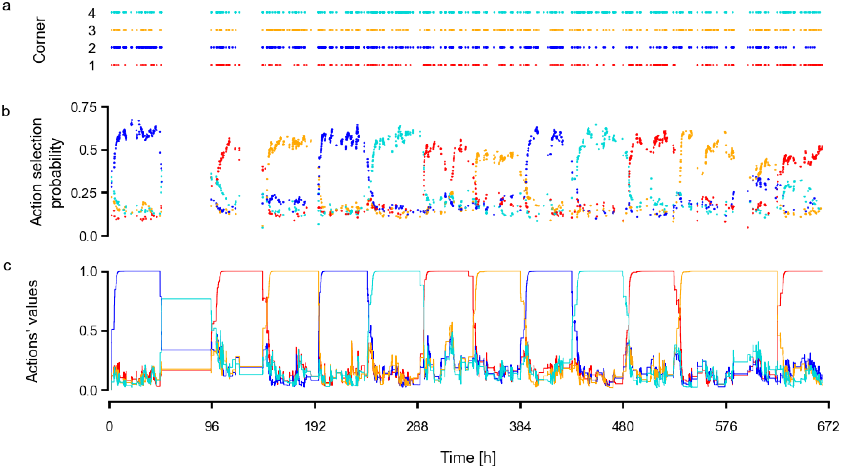
Complete behavioral data from all the experimental phases for mouse 14. **a** raster plot of mouse activity. Every dot represents entry into a given color-coded corner. **b** valuations learned by the mouse. **c**, soft-max probabilities to enter the corner given the valuation. Note that only the self-learning part is shown here.

To compare the adequacy of the different models in the description of behavior of each mouse we computed the Bayesian (Schwarz) Information Criterion for the models tested ^30^. This is the negative log-likelihood that the observed data were generated by the given model (*L*) penalized by the number of parameters *k* and data points *n*: *BIC* = *k* ln *n* − 2 ln(*L*). Supplementary Table 2 gives BIC values for every model and each mouse. To facilitate understanding of these results, in Fig. 4**a** we show these BIC values normalized to the unit interval. To normalize for each mouse *i* and model *m* we subtract the minimum BIC over models and normalize the result by the difference between the maximum and minimum:

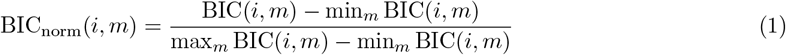

**Figure 4:**
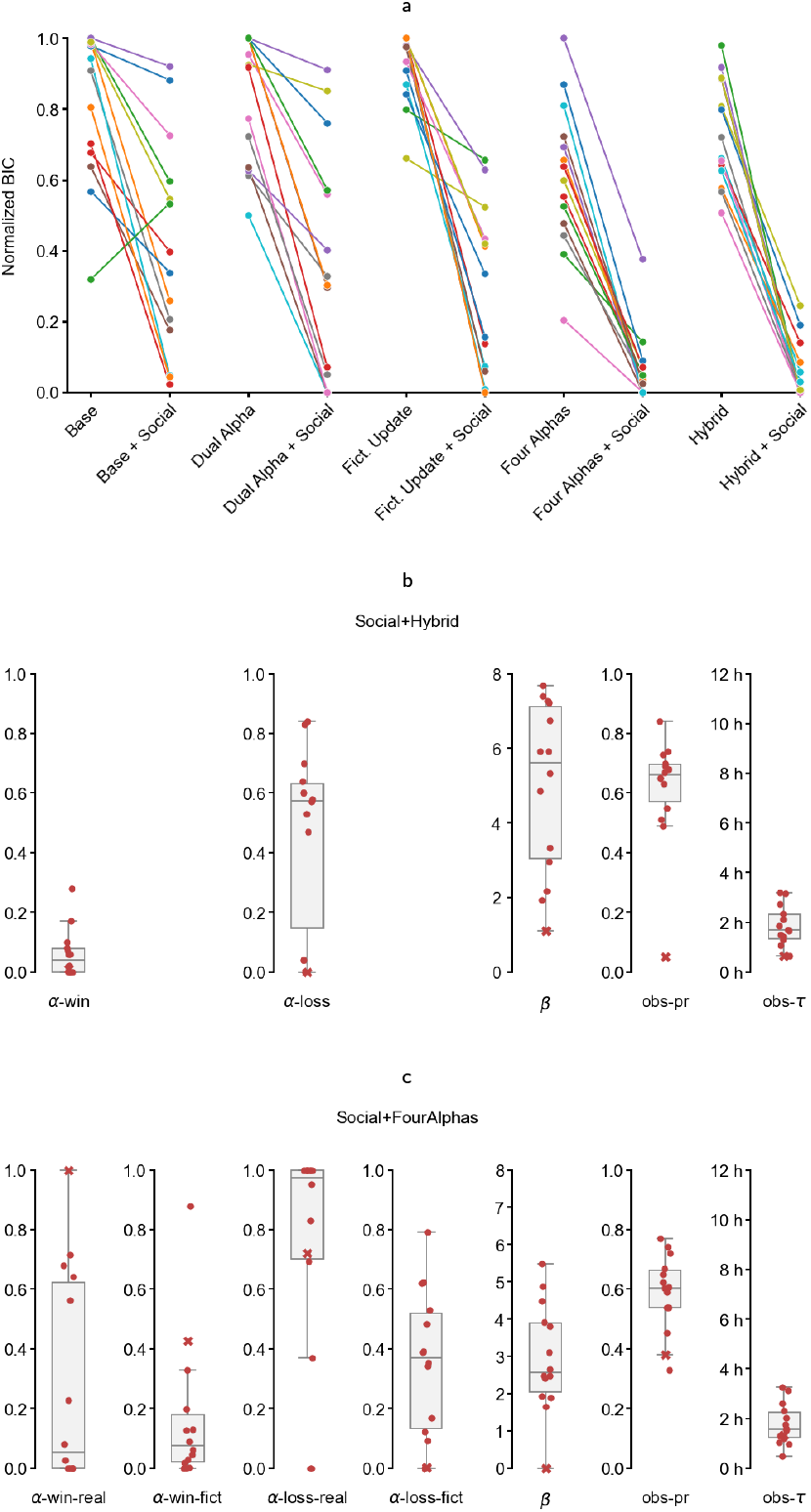
**a** Bayesian Information Criteria for every model fit and every mouse minus minimum BIC for every mouse. **b** The distribution of parameter values for the Hybrid+Social model. **c** FourAlphas+Social model fitted to the behavior of all the mice.

In this way, the most parsimonious model for the given mouse, which has the smallest BIC, is normalized to 0, while the model with the largest BIC is normalized to 1. Interestingly, we can see that for every mouse the best model always requires accounting for the social aspects. Also, the basic RW-learning, even when the social effects are taken into account, is never the best explainer of the obtained data.

We pair the results for every model type for self-learning variant with the variant including social effects. It is apparent that in every case the addition of the social effects significantly improves explainability of the model. It is also apparent that Hybrid+Social and FourAlphas+Social are performing best. The model which explains the data best overall is Hybrid+Social and we present the distribution of its parameters in Fig. 4**b**. This was the best model for 6 animals but also the summary BIC is the smallest among all the models, Table 2. The second best is the FourAlpha+Social model which disambiguates true learning from inverse fictitious learning at the cost of extra two parameters which occasionally leads to stronger penalization but still for 4 animals this is the best model. Fig. 4**c** shows the parameters for the FourAlpha+Social model fitted to the observed behavior of all mice. The first thing we observe is that the alpha-win parameter is close to 0 for most mice. This means that the valuations are modified only slightly when the animal receives reward. On the other hand, in most cases alpha-loss is around 0.6, which means that the learning occurs mostly during losses.

The next striking observation is that the probability for a mouse to make choice based on the social signal is about 0.65. Thus, in two cases out of three the mice choose the socially preferred corner over what they learned individually. One exception we see in this plot is the value for mouse 13. The performance of mouse 13 was the worst in terms of reward consumption. It seems it did not learn well but also the fits of our models to this mouse behavior were not reliable, so we cannot discuss this mouse in a meaningful way. A more reliable estimates of this mouse behavior were obtained in our complementary approach ^31^. In any case, in any model fitted to this mouse, the social probability parameter was among the lowest, consistent with relatively poor performance of this animal. Finally, the observation time in which we assume the mice build their model of social preference of corners comes around 2 hours which seems realistic.

## Discussion

### Summary

Making precise statements about biological processes, whether in biophysics or behavior, requires both well-controlled experiments and precise conceptual and theoretical frameworks which can facilitate description of data and meaningful analysis. Here, we proposed a general framework to describe and analyze behavior of animal groups where each animal’s behavior is described by a series of events, discrete in time. It is then practical and convenient to consider this behavior in terms of transitions between discrete states. The example experimental setup for which we presented our framework explicitly is that of the Intellicage system. Here, a mice cohort of up to 16 mice, moves in a cage with four corners and behavior of each mouse can be quantized as being in one of the four corners or outside.

In this framework the behavior of animals is described as a marked inhomogeneous point process where the animals can assume different states of their choosing (that is, enter different corners) according to the valuations attached to those states. Those valuations change with time due to individual or group learning. In the simple example discussed here each animal can assume one of five states: be in one of four cage corners or outside. The probability to visit any corner changes in time as the animal explores rewards available in different corners as well as when it observes its mates’ behavior. We considered five different learning models with two variants each, assuming self-learning only or including a social effect of the peer behavior.

Our results place significant quantitative constraints on any possible neural implementation of individual and social decision making. For example, we showed that the social hybrid model or the social four alpha model are the most likely to explain the observed behavior. This indicates that the simplest RW-learning models are insufficient and more sophisticated neural implementation is required to accommodate both real and fictitious learning with different speeds for rewards and no rewards.

### Implications

The simple framework of the Intellicage allowed us to make several interesting observations. Most strikingly, for every animal in the cohort, the models assuming self-learning combined with reliance on the mean-field signal from the rest of cohort were always preferred to models based on individual learning only, Fig. 4 **a**. Depending on animal different models were selected by the BIC, with the Hybrid+Social model being the most common choice (6 animals), followed by FourAlpha+Social (4 animals), DualAlpha+Social (3 animals) and FictitiousUpdate (1 animal — mouse 13, with inconsistent fits, see results).

We found it surprising that, on average, in two cases out of three the animals made a choice based on the mean behavior of the cohort rather than on self-learned corner valuations. This strong reliance on mean behavior signal led to relatively uniform probabilities of corner choice. In consequence, the behavior had strongly exploratory character with limited reward exploitation. Nevertheless, the likelihood analysis led to very specific model selection and one could clearly observe that despite significant exploration, learning the reward corner took place within around 10 hours.

Interestingly, the simplest RW-learning was never implied. Therefore, since more complicated models were systematically indicated, the postulated or equivalent models must be somehow implemented in the brain. This leads to natural questions about the where and how can this be implemented. The proposed approach allows to describe mice cohorts but also individuals within cohorts as it is possible to fit individualized models to individual animal data within a cohort. This allows to study group coherence and detect outliers, as exemplified here by animal 13 which had difficulty learning.

### Possible extensions

We chose the Intellicage system because although several different cage designs have been in use in group studies ^16–20;24^ most of them have been developed by individual labs making comparisons between results coming from different groups difficult. The Intellicage remains one of the few commercially available products with broad community adoption (at least 80 groups, 3500 animals tested as of 2020^27^). Our presentation here is focused on this design to have direct appeal in the community but our computational framework can be adopted and generalized also to other cages where behavior is monitored by discrete events in time ^18–20^.

In presenting the proposed analytical framework we decided to focus on a relatively simple case where we consider only visits to corners. This could easily be extended by considering more complex definition of events. For example, in the Intellicage, we could add further indices indicating interactions with specific bottles within corners or licking status. Combining the point process approach with learned valuation-based decision making could be extended to other cases of animal populations exploring a common space where the behavior is tracked by discrete events happening in discrete times, such as in Eco-HAB system ^20;22^ or in more naturalistic contexts where the population explores an arbitrary arena or a maze and interacts with discrete elements changing their states, such as AutonoMouse ^32^. It could also be adapted for more involved automated home cage monitoring systems, combining RFID gating with video monitoring, beambreaks, lickometers, air/pressure or capacitive floor sensors, and embedded operant modules to time-stamp discrete events while preserving group housing and minimizing handler effects^33–39^. Note that after video preprocessing using deep learning tools such as the Deep Lab Cut^15;40^ and translation into time-stamped events, the marked point process approach might be a useful framework for further conceptual processing.

The experimental example we used to illustrate the proposed formalism and technique uses appetitive learning but our approach could also be applied in the aversive setting or for addiction.

### Experimental design and possible extensions

In designing the experiment we considered an optimal division into the minority and majority groups. Any choice here is a trade-off between enhancing the desired effects or improving statistics of the minority group. Clearly, the choice we made of minority group of 2 individuals and majority group of 10 improves the chances of observing the effect, as the majority is 6 times bigger then the minority and can strongly pull minority towards majority preferred corner. This is at the price of poorer statistics for the minority group. We discarded the case of minority group of 1 which could possibly be of interest to study as here all social cues would be misleading. As we reduced the difference between sizes of majority and minority groups so that both equaled 7 we would obviously expect a general confusion as the effect of “minority” of 7 would be bigger than that of “majority” of 6 (excluding self). It could be of interest to check if in that case the mice are able to discover the identity of its team in the course of time. Given that our model is generative that is it allows to simulate the process one could make predictions for arbitrary variants of our experiment, varying the composition and size of majority and minority groups at will and making predictions to further improve the modeling when significant deviation between said predictions and obtained results is observed.

### More complex estimation

Our estimation here assumes independence of transition dynamics and learning. This may be too simplistic. Further, not every visit is associated with drinking. So we have really two types of visits: those related to drinking (goal-oriented) and those reflecting more exploratory behavior (idle mode) which we explore in our companion paper ^31^. There we also go beyond the mean-field approach, conditioning behavior on individual animal contributions which allows us to identify social structure within the cohort (see also ^22^).

We assumed here that the strategy used by an animal is fixed and the behavior changes as a consequence of learning which affects the learned valuations but not the strategy. In principle one can imagine that the strategy itself may change with time. For example, if the animal is able to detect while in minority that its reliance on the group signal leads to a decrease in the success rate the animal might reduce this reliance on social signal. However, the amount of data needed for reliable parameter estimation is relatively large and in consequence estimates on data fragments, specifically individual experimental phases, incur substantial errors. One can compromise by considering estimation in phases, which is also discussed in ^31^. One can also imagine estimation strategies assuming slow parameter variation in time which might mitigate the estimation problems. This will be explored in the future.

### Outlook

The presented research leaves more questions opened than answered. One may wonder if the strategy to follow the crowd was preferred by the mice natively or if they learned it to best cope in the situation they found themselves. One may ask if a mouse can learn to completely ignore the social cues. Such a strategy would be optimal in a situation where the social cues would be decorrelated from individual cues for a given mouse. To check it we could do an experiment where we select 1 or 2 mice and their reward placement is selected randomly with respect to randomly selected placement of reward for the rest of the mice. Another interesting question would be the effect of reward corner shift at different times of the circadian cycle. Does it matter for the mouse if we shift the position of the reward in the middle of the active / inactive phase? It seems that value learning takes about 12 hours. Do the strategies change if we shift reward faster? Say, every 8 hours, or in randomized intervals? The Intellicage systems allows to pose these and related questions and answer them using the presented analytical framework. We hope to report on some of them in the future.

## Acknowledgments

BJ, ML, and DKW acknowledge support by the project POIR.04.04.00-00-14DE/18-00 carried out within the Team-NET programme of the Foundation for Polish Science co-financed by the European Union under the European Regional Development Fund. The authors declare no conflict of interest.

## Methods

### Experimental methods

Experiments were performed on 14 female C57BL/6 mice from the colony bred at the Maj Institute of Pharmacology of the Polish Academy of Sciences in Kraków. Mice were housed in Plexiglas cages (Type II L, 2–5 animals per cage) with aspen laboratory bedding (MIDI LTE E-002, Abedd) and nest-building material, under a 12 hr light-dark cycle, with an ambient temperature of (22 ± 2)°C. Animals had ad libitum access to water and chow (RM1 A (P), Special Diets Services) and were provided with a piece of aspen wood for chewing. Mice had 11 to 15 weeks of age and weighed 19.3 to 22.3 g at the start of the experiments. Mice were implanted with radio frequency identification chips (RFID chips, UNO PICO ID, AnimaLab, Poland) before being introduced to the IntelliCage. All experiments were conducted in accordance with the European Union guidelines for the care and use of laboratory animals (2010/63/EU). Experimental protocols were reviewed and approved by the II Local Bioethics Committee in Krakow (permit LKE 109/2021).

Behavior analysis was conducted using an IntelliCage apparatus (New Behavior, Switzerland) ^41^. The IntelliCage consists of a base transparent plastic box (55 × 37.5 × 20.5 cm) with a metal cover and custom corner compartments. Each of the cage corners is a small chamber that houses two 250 ml bottles, with nozzles accessible through guillotine doors, Fig. 1**a**. The size of the corner allows only one animal to enter the corner and access the bottles. Antennas inside the corners detect the RFID tags and report the number of the tag to the IntelliCage Controller software (TSE, Germany), which triggers programmed events. The cage recorded the following parameters: temperature and luminosity, presence of an animal in a corner, the crossing of photocell beams placed in the doors leading to the bottles, and lickometer contacts (the animal closing a circuit between the floor grating and the metal dipper of the bottle).

The behavioral procedure started with adaptation followed by the main procedure with a choice between water and saccharin bottles. The experimental schedule is provided in Fig. 1**c–e**. When introduced to the cage mice were allowed free access to all water bottles in all corners with all doors fixed open for 4 days (not shown). The adaptation to the basic experimental condition with one door open per corner lasted 4 days with a change of accessible bottles in every corner after two days. Next, the corner doors became closed by default, and when an animal entered a corner and remained there for 1 s, one of the doors would open providing access to a bottle. The door would close 10 s after licking started or when the animal left the corner, Fig. 1**c**. In the final adaptation stage, one of the accessible bottles contained 0.1 % saccharin solution. The position of the saccharin solution was changed every 24 hours for 8 consecutive days, Fig. 1**d**. Therefore, adaptation to the experimental environment lasted 16 days in total.

During the main experimental procedure, animals again had access to 3 water bottles and 1 saccharin solution bottle, Fig. 1**e**, however, they were split into two groups, Fig. 1**b**. 12 animals were assigned to ‘majority’ and 2 to ‘minority’ groups. The two groups differed in the assigned access to the saccharin solution bottle, which was provided in different corners. Thus, if social cues played a role in learning the placement of the saccharin bottle, then ‘minority’ animals would be misdirected with regard to the position of the reward. The assignment of groups changed every 4 days, in a way that every animal was placed in the minority group for one 4-day period.

### Formal description of mouse behavior in Intellicages

We propose to describe mice behavior in the Intellicages in the language of marked point processes ^28^. Focusing on the discrete transitions of each mouse state which are entering and leaving a corner, they are like spikes sent by neurons signaling discrete events. Therefore, we can represent the behavior of the whole cohort with a rasterplot where every row represents a mouse and every event represents a transition with color representing the corner number or the arena. We call this approach the kinematics of mice and the rasterplot “colored spikes”. Our construction below follows standard description of spiking ^42–45^.

Assume *N* mice in a cage and an experiment of duration *T* . We say a mouse *i* is in a state *k* = 1 to 4 if it entered corner *k*. Otherwise we say it is outside or in the “arena” and has a state 0. Thus we have a discrete state space of *K* = 5 states. Note an exclusion principle: only one mouse at a time can be in any nonzero state.

Given available information, complete behavior of every mouse is described by its state as a function of time *s*_*i*_(*t*) (trajectory, Fig. 5.A): ∀ _*t*∈[0,*T*]_ *s*_*i*_(*t*) ∈ {0, 1, 2, 3, 4 }. Equivalent information is contained in the times *t*_*i,j*_ at which the mouse *i* changes its state to *s*_*i,j*_ = *k* (enters corner *k* or leaves a corner if *k* = 0). Then the complete description of the mouse behavior is given by the sequence of times and states taken,

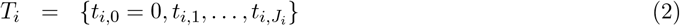

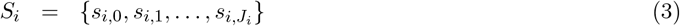

where *t*_*i,j*_ ∈ [0, *T*] is the time of state change, *s*_*i,j*_ is the new state, and the *J*_*i*_ is the number of state changes (transitions). Note that in this description when the state is either a corner or the arena, *J*_*i*_ is typically twice the number of mouse visits in the standard Intellicage terminology: we treat the beginning and the end of a visit as individual events. We assume *s*_*i*,0_ is the initial state of the mouse which is 0 if *t* = 0 coincides with the true beginning of the experiment.

**Figure 5:**
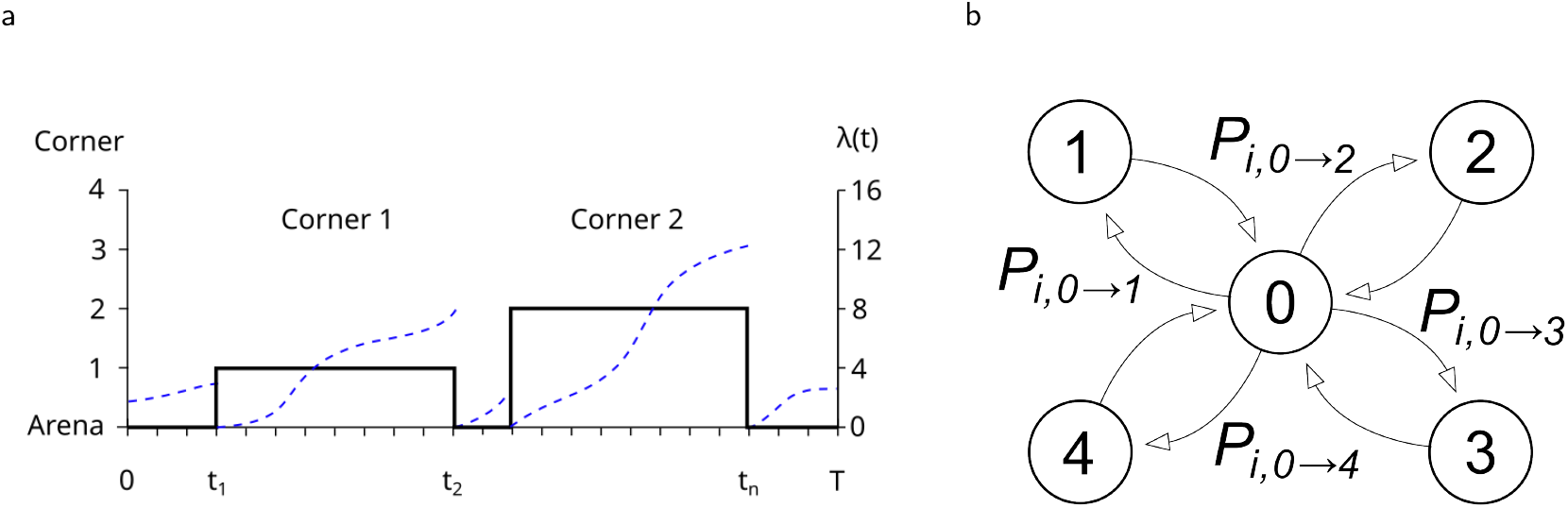
**a** Example trajectory of a mouse. The mouse is in the arena during *t* ∈ [0, *t*_1_) ∪ (*t*_2_, *t*_3_) ∪ (*t*_*n*_, *T*). It is in corner 1 during *t* ∈ (*t*_1_, *t*_2_) and in corner 2 during *t* ∈ (*t*_3_, *t*_*n*_). We have *s*_0_ = *s*_2_ = *s*_*n*_ = 0, *s*_1_ = 3, *s*_3_ = 4. Dashed blue line is the hazard function *λ*(*t*) for the mouse. See text for details. **b** Transition probabilities between arena and the corners. From a corner you can move only to the arena. From the arena you can move to one of four corners.

To describe the dynamics we introduce counting functions *N*_*i,j*→*k*_(*t*) which count the number of transitions of mouse *i* from state *j* to *k* up to time *t*. Note that

1. for *j, k* ≠ 0 we have *N*_*i,j*→*k*_(*t*) = 0,
2. *N*_*i,j*→*j*_(*t*) = 0.

This is a consequence of the construction of the IntelliCage and reflects the fact that a mouse in the arena can go to any corner but a mouse in a corner can only move out to the arena.

We further introduce the history of mice behavior up to time *t* as

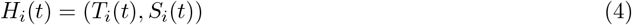

where

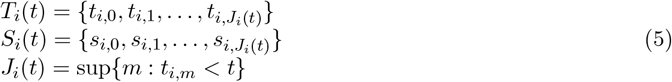

and the overall history of the population

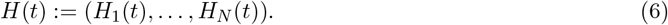

We can also encode the trajectory as

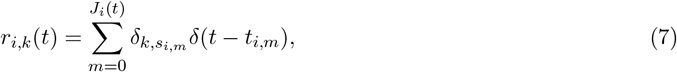

using Kronecker and Dirac delta notation. This description is similar to the typical description of spiking in neural networks^43^. We call it a “colored spike train” or a “marked spike train”, as each “spike” (event) carries a “mark” which is the state we enter.

### Stochastic models of mice behavior

Given apparent randomness of mice behavior we resort to probabilistic modeling and interpret mice trajectories as realizations of a stochastic process. We postulate here that the behavior of animals making spontaneous decisions in an environment offering discrete choice options can be described, analyzed and modeled within the framework of marked point processes. Here, we show how this framework can be applied to mice groups housed in the Intellicage.

We consider the process governing the mice behavior to be a marked point processes ^28^. For each mouse the transition times can be considered a realization of a simple point process ^44;45^, however, each event carries a mark which is the state entered.

We describe the combined dynamics through stochastic intensity also called a hazard function ^42;44;45^:

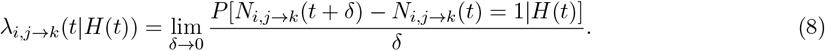

It is the momentary event rate for mouse *i*, given the history of all mice *H*(*t*) up to time *t*, of making a transition from state *j* to *k*. This is the fundamental quantity in the theory needed for simulation, analysis and model estimation and is the most general formulation we consider here. To reach a level when this becomes amenable to estimation we will restrict our attention to specific classes of models.

We first impose a product structure on *λ*:

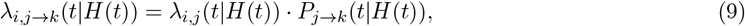

where the first factor

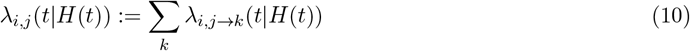

is the hazard function generating a transition of mouse *i* which is at time *t* in state *j*, in other words, *λ*_*i,j*→*k*_(*t* |*H*(*t*)*dt* is the probability that a state change will take place in the infinitesimal interval (*t, t* + *dt*), given the history *H*(*t*). The second factor is the conditional probability of a transition from state *j* to *k* at time *t* given the history, which includes possible interactions with the cage reflected through a learning process.

In our setting, transitions may only take place from arena, *j* = 0, to one of the corners, *k >* 0, given by *λ*_*i*,0→*k*_, or from a corner to the arena, *λ*_*i,k*→0_. We now assume that both are modulated by a global activity function, 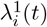, predominantly reflecting the daily activity cycle. Further, we assume that the chances to change state depend mainly on the time since moving into the present state, *τ* = *t* − *t*^∗^, where *t*^∗^ is the time of transition into the present state, but these chances will differ between the arena and the corners. Assuming different probabilities for different corners might take into account biases but given available data is difficult to estimate. However, the different statistics of visit durations between rewarded and unrewarded corners is accounted for by including *λ*^3^ factor to modulate transition chances in unrewarded corners and *λ*^4^ in the rewarded ones.

This leads to the following formulation:

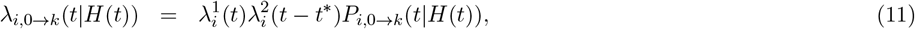

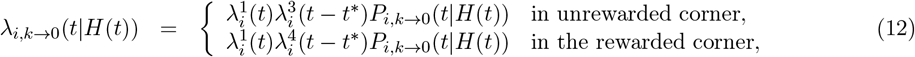

where reward is sacharine provided in one of the bottles.

Since from any corner *k* we can only leave into the arena, we require *P*_*i,k*→0_(*t* |*H*(*t*)) ≡1 which leads to the final form of our model

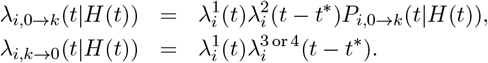

Observe that the form of the hazard function we adopt is the Inhomogeneous Markov Interval model proposed by Kass and Ventura for spiking activity ^46–48^. Since this is a marked point process we call it the Marked Inhomogeneous Markov Interval model (MIMI).

### Individual learning

We assume that the conditional probability for a mouse to enter corner *k* is a function of the value it attributes to the corner, *Q*_*i,k*_(*t*). We consider general, abstract value-learning models, derived from Rescorla-Wagner rule of choice valuation, all different relatives of basic Q-learning^1^. We assume mouse *i* keeps its valuation of corner *k, Q*_*i,k*_(*t*). We use soft-max rule for the transition probability:

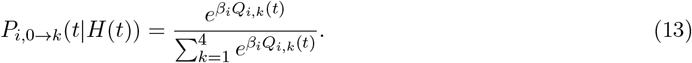

Valuations are updated when leaving a corner according to one of reinforcement learning models ^1^. We consider five individual learning model variants^29;49–51^ described in Table 1.

**Table 1:** Individual learning models. *a* — chosen corner, *b* — any other corner.

| Model name | parameters | model equations |
| --- | --- | --- |
| Basic | $\alpha, \beta$ | $Q_{a,n+1} = Q_{a,n} + \alpha(R_n - Q_{a,n})$<br>$Q_{b,n+1} = Q_{b,n}$ |
| Dual Alpha | $\alpha_{win}, \alpha_{loss}, \beta$ | $Q_{a,n+1} = Q_{a,n} + \alpha_{win}(R_n - Q_{a,n})\vert _{R_n=1}$<br>$Q_{a,n+1} = Q_{a,n} + \alpha_{loss}(R_n - Q_{a,n})\vert _{R_n=0}$<br>$Q_{b,n+1} = Q_{b,n}$ |
| Fictitious Update | $\alpha, \beta$ | $Q_{a,n+1} = Q_{a,n} + \alpha(R_n - Q_{a,n})$<br>$Q_{b,n+1} = Q_{b,n} + \alpha((1 - R_n) - Q_{b,n})$ |
| Hybrid | $\alpha_+, \alpha_-, \beta$ | $Q_{a,n+1} = Q_{a,n} + \alpha_+(R_n - Q_{a,n})\vert _{R_n=1}$<br>$Q_{a,n+1} = Q_{a,n} + \alpha_-(R_n - Q_{a,n})\vert _{R_n=0}$<br>$Q_{b,n+1} = Q_{b,n} + \alpha_+(1 - R_n - Q_{b,n})\vert _{R_n=1}$<br>$Q_{b,n+1} = Q_{b,n} + \alpha_-(1 - R_n - Q_{b,n})\vert _{R_n=0}$ |
| Four Alpha | $\alpha_{win}, \alpha_{loss}, \alpha_{fict.win}, \alpha_{fict.loss}, \beta$ | $Q_{a,n+1} = Q_{a,n} + \alpha_{win}(R_n - Q_{a,n})\vert _{R_n=1}$<br>$Q_{a,n+1} = Q_{a,n} + \alpha_{loss}(R_n - Q_{a,n})\vert _{R_n=0}$<br>$Q_{b,n+1} = Q_{b,n} + \alpha_{fict.win}(1 - R_n - Q_{b,n})\vert _{R_n=1}$<br>$Q_{b,n+1} = Q_{b,n} + \alpha_{fict.loss}(1 - R_n - Q_{b,n})\vert _{R_n=0}$ |

**Table 2:**
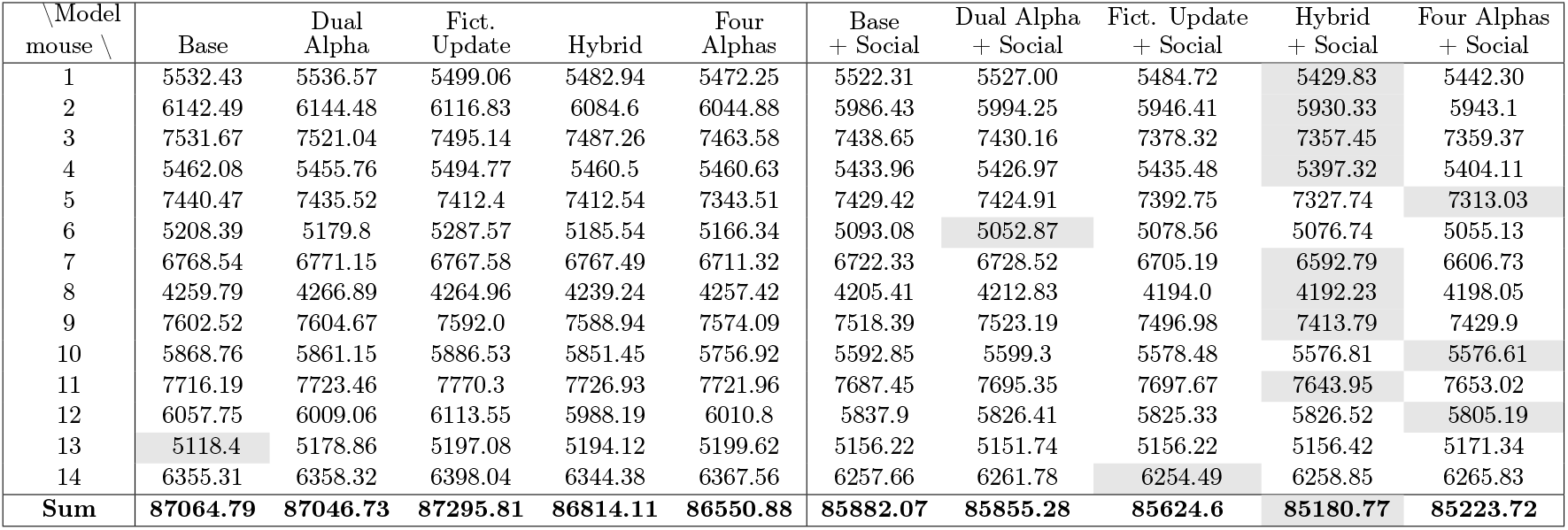
Bayesian Information Criteria for model fits to experimental data. Rows: mouse number. Column: model. Highlighted are the minima for a given mouse indicating the model selected by the BIC.

| \Model<br>mouse \ | Base | Dual<br>Alpha | Fict.<br>Update | Hybrid | Four<br>Alphas | Base<br>+ Social | Dual Alpha<br>+ Social | Fict. Update<br>+ Social | Hybrid<br>+ Social | Four Alphas<br>+ Social |
| --- | --- | --- | --- | --- | --- | --- | --- | --- | --- | --- |
| 1 | 5532.43 | 5536.57 | 5499.06 | 5482.94 | 5472.25 | 5522.31 | 5527.00 | 5484.72 | 5429.83 | 5442.30 |
| 2 | 6142.49 | 6144.48 | 6116.83 | 6084.6 | 6044.88 | 5986.43 | 5994.25 | 5946.41 | 5930.33 | 5943.1 |
| 3 | 7531.67 | 7521.04 | 7495.14 | 7487.26 | 7463.58 | 7438.65 | 7430.16 | 7378.32 | 7357.45 | 7359.37 |
| 4 | 5462.08 | 5455.76 | 5494.77 | 5460.5 | 5460.63 | 5433.96 | 5426.97 | 5435.48 | 5397.32 | 5404.11 |
| 5 | 7440.47 | 7435.52 | 7412.4 | 7412.54 | 7343.51 | 7429.42 | 7424.91 | 7392.75 | 7327.74 | 7313.03 |
| 6 | 5208.39 | 5179.8 | 5287.57 | 5185.54 | 5166.34 | 5093.08 | 5052.87 | 5078.56 | 5076.74 | 5055.13 |
| 7 | 6768.54 | 6771.15 | 6767.58 | 6767.49 | 6711.32 | 6722.33 | 6728.52 | 6705.19 | 6592.79 | 6606.73 |
| 8 | 4259.79 | 4266.89 | 4264.96 | 4239.24 | 4257.42 | 4205.41 | 4212.83 | 4194.0 | 4192.23 | 4198.05 |
| 9 | 7602.52 | 7604.67 | 7592.0 | 7588.94 | 7574.09 | 7518.39 | 7523.19 | 7496.98 | 7413.79 | 7429.9 |
| 10 | 5868.76 | 5861.15 | 5886.53 | 5851.45 | 5756.92 | 5592.85 | 5599.3 | 5578.48 | 5576.81 | 5576.61 |
| 11 | 7716.19 | 7723.46 | 7770.3 | 7726.93 | 7721.96 | 7687.45 | 7695.35 | 7697.67 | 7643.95 | 7653.02 |
| 12 | 6057.75 | 6009.06 | 6113.55 | 5988.19 | 6010.8 | 5837.9 | 5826.41 | 5825.33 | 5826.52 | 5805.19 |
| 13 | 5118.4 | 5178.86 | 5197.08 | 5194.12 | 5199.62 | 5156.22 | 5151.74 | 5156.22 | 5156.42 | 5171.34 |
| 14 | 6355.31 | 6358.32 | 6398.04 | 6344.38 | 6367.56 | 6257.66 | 6261.78 | 6254.49 | 6258.85 | 6265.83 |
| Sum | 87064.79 | 87046.73 | 87295.81 | 86814.11 | 86550.88 | 85882.07 | 85855.28 | 85624.6 | 85180.77 | 85223.72 |

**Table 3:** Bayesian Information Criteria for model fits to simulated data of 14 independent “mice” learning reward with Q-learning. Row: mouse number. Column: model. Highlighted are the minima for a given mouse indicating the model selected by the BIC.

| \Model<br>mouse \ | Base | Dual<br>Alpha | Fict.<br>Update | Hybrid | Base<br>+ Social | Dual Alpha<br>+ Social | Fict. Update<br>+ Social | Hybrid<br>+ Social |
| --- | --- | --- | --- | --- | --- | --- | --- | --- |
| 1 | 3623.71 | 3630.82 | 3633.26 | 3631.32 | 3632.84 | 3639.89 | 3638.91 | 3640.53 |
| 2 | 3763.77 | 3769.09 | 3766.82 | 3771.90 | 3778.29 | 3783.60 | 3781.33 | 3786.41 |
| 3 | 3832.62 | 3838.76 | 3845.42 | 3835.17 | 3846.00 | 3852.47 | 3857.11 | 3849.13 |
| 4 | 3600.86 | 3607.71 | 3608.50 | 3610.67 | 3614.10 | 3620.94 | 3619.73 | 3623.88 |
| 5 | 3718.42 | 3725.52 | 3732.08 | 3731.41 | 3732.94 | 3739.78 | 3746.60 | 3745.61 |
| 6 | 3693.10 | 3700.22 | 3699.17 | 3702.19 | 3706.14 | 3713.17 | 3711.63 | 3715.22 |
| 7 | 3716.44 | 3723.41 | 3725.67 | 3725.93 | 3729.37 | 3736.29 | 3736.80 | 3738.66 |
| 8 | 3683.87 | 3690.48 | 3700.76 | 3692.16 | 3691.87 | 3698.57 | 3701.21 | 3700.61 |
| 9 | 3562.03 | 3569.05 | 3562.95 | 3569.64 | 3575.81 | 3582.83 | 3576.18 | 3583.21 |
| 10 | 3717.63 | 3724.17 | 3728.08 | 3728.28 | 3730.59 | 3734.47 | 3740.08 | 3741.17 |
| 11 | 3713.82 | 3720.92 | 3718.30 | 3724.53 | 3728.32 | 3735.29 | 3732.80 | 3739.03 |
| 12 | 3864.61 | 3871.75 | 3868.86 | 3873.87 | 3879.19 | 3886.33 | 3883.38 | 3888.45 |
| 13 | 3746.92 | 3753.59 | 3744.35 | 3750.71 | 3761.28 | 3767.97 | 3758.26 | 3763.77 |
| 14 | 3786.91 | 3793.71 | 3799.49 | 3797.65 | 3794.79 | 3801.90 | 3801.74 | 3804.58 |
| Sum | 52024.69 | 52119.17 | 52133.72 | 52145.43 | 52201.51 | 52293.50 | 52285.74 | 52320.26 |

### Social aspects of learning

We consider two types of models representing individual and social learning. In individual learning models we assume that the mouse makes decisions based on its own experience according to one of the Q-learning variants described above. In the social learning models we consider the mouse selects a corner depending on the history of visits of all the other mice with probability *γ*, and with probability 1 − *γ* it makes a decision based on its own experience.

The resulting transition probability is given by

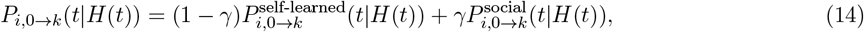

where *P* ^self-learned^ is the soft-max probability for one of the individual learning models, Eq. (13), and

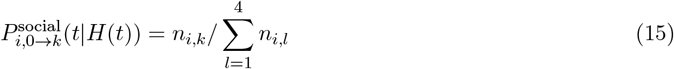

is the time weighted estimation of the popularity of a given corner. Here

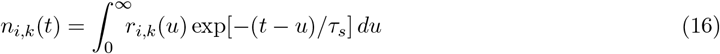

where *τ*_*s*_ quantifies the time span of the social observation and *r*_*i,k*_(*t*) is the collection of visits of mouse *i* in corner *k* given by Eq. (7).

### Estimation

The probability of observing the mouse trajectory *H*(*T*) (Fig. 5) under the assumption of specific model is given by

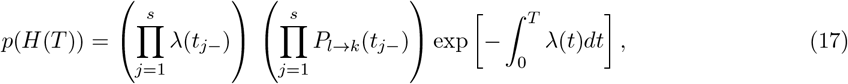

where *λ*(*t*_*j*−_) is the limit of *λ* at *t* from the left, etc. Thus its log-likelihood is

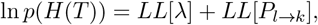

where

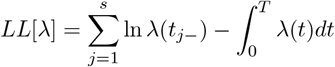

and

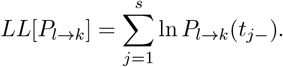

Note that under the assumptions we have made here these two terms depend on different variables and so can be optimized independently. Specifically, the first term depends only on the assumed models of the hazard function, which is a product of spline-based *λ*^1^ and similarly modeled *λ*^2,3,4^. The second term depends on the assumed model of learning (individual or social).

Fig. 6 shows example results of fitting the Basic model (simple Q-learning) to the behavior of mouse 7. The fragment shows four days of the experimental phase E4. Panel **a** shows in blue kernel interpolated intensity of transitions, which corresponds to the firing rate of spiking data ^43^, red line shows the Poissonina parts of the estimated intensity 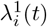. Panel **b** shows corner visits. Panel **d** shows estimated corner valuation *Q*_7,*k*_(*t*) using the base model. We can see that according to the model this mouse learns the location of the reward very fast: it takes about 10 hours for the mouse to attribute maximum of 1 to the rewarded corner, but also the same time to forget it and learn the new one. Interestingly, due to the soft-max rule the difference between the preferred corner and the rest as shown in panel **c** is rather limited but the model can still discern it. The maximum probability *P*_*i*,0→*k*_ is around 0.4, about twice as much as for the remaining corners thus leaving substantial space for exploration of all corners.

**Figure 6:**
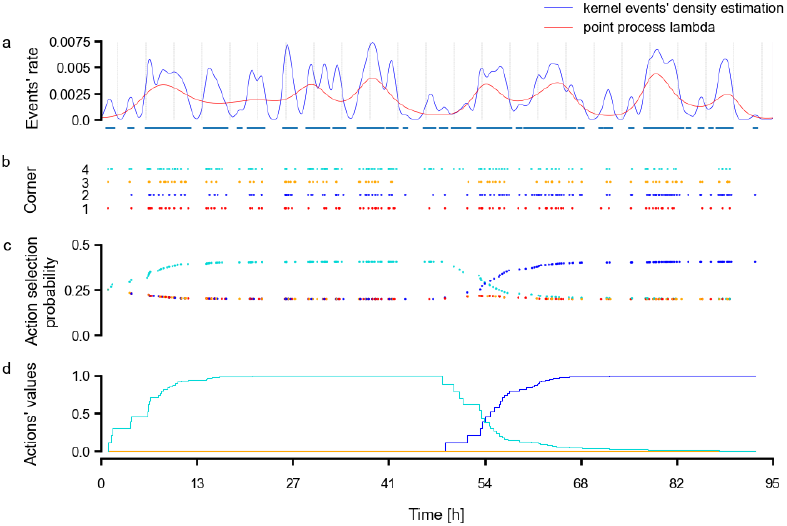
Example fit of the Base model for mouse 7 in the experimental phase E4. **a** blue: kernel interpolated intensity of transitions. Short vertical bars on the lower horizontal axis: individual transitions; red: estimated intensity 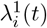 (the Poissonian part). **b** corner visits. **c** probabilities to enter a corner computed from the fit in panel **d**) with soft-max rule. **d** estimated corner valuation *Q*_14,*k*_(*t*) in the base model.

### Technical aspects of estimation

We fit each animal’s behavior independently. The models consist of two parts, the stochastic intensity function built from *λ*^1,2,3,4^ and the learning part implemented as one of the reinforcement learning models. The factors of the intensity function are constructed as exponentials of spline functions with knots set every few hours (3 hours by default) for *λ*^1^, and with 20 s for *λ*^2,3,4^. We set periodic boundary conditions on data by cyclic data padding, ensuring smooth fit at boundaries. This facilitates modeling of a periodic process, such as a circadian rhythm, despite using data from a restricted time interval. We use exponential to ensure positivity of the intensity, effectively implementing a generalized logistic model. Estimation was performed in Python with scipy.interpolate.splev function for cubic splines. For fitting and simulation of the point process the time was discretized with *dt* = 1*s*.

Learning reward placement was modeled with reinforcement learning of the valuations of four corners. Initial values of corner valuations *Q*_*i,n*_, before the adaptation phase, were all set to 0. This arbitrary setting can be justified by the relatively long duration of the experiment (27 days), with many alternations between corners, so the choice of these initial values should not introduce a significant bias in the results. First, we fit the model to the initial adaptation phase, where reward was in one and the same corner for all animals, and then take the final corner-values as initial values of a model at the start of the first experimental phase. Similarly, when fitting individual phases, we propagated corner valuations learned in the preceding phases. We set reward (saccharine) value to 1, no reward (water) to 0. Setting reward to some other value would result in rescaling the values of *α*.

In estimation of models with social interactions from experimental data, to reduce computational demands we used the mean-field social signal representing a weighted average of visits of every animal to each corner. This was computed by taking the time of each visit of every mouse *t*_*s*_, for each corner separately, within a 48 h window preceding a given time *t*, contributing to the mean-field average visit probability with weight representing exponential decay with time scale *τ*_*s*_ which was also fitted from data (Eq. 15). Rather then compute a different signal for every observer from visits of everybody else we included visits of all 14 mice for each observing mouse in this mean-field signal thus always including the visits of the observer. This simplified computations but could introduce on average a 1:13-fold bias. Note that during simulations of the social influence in the social signal (for validation of methods, see next section) we included only visits of the other 13 observed mice, but nonetheless we were able to recover the true underlying model.

Models were fit by minimizing negative log-likelihood (nll) of the data under a given model using Python scipy.optimize.minimize function, separately for the point process and learning parts. We used either Nelder-Mead simplex algorithm, or, especially for the point process part, L-BFGS-B algorithm (using gradient information computed numerically with finite differences method). For problems with constraints (*λ*^2,3,4^ modeled with splines) we used trust-constr algorithm, with nonlinear constraints on the splines’ parameters demanding that the renewal factors in stochastic intensity *λ* ≡ *λ*^2,3,4^ satisfy: (1) *λ*(*τ* = 0) = 0, (2) *λ*(*τ* = 120*s*) = 1, (3) *λ*^*′*^(*τ* = 0) = 0, (4) *λ*^*′*^(*τ* = 120*s*) = 0, (5) 0 *< λ*(*τ*) *<* 1. When using the simplex method the data nll was set to return Inf on the boundary of the parameter space. Parameter bounds for the optimization procedures were given as follows: *α*_*i*_ ∈ (0; 1); *β* ∈ (0; 25); *γ* ∈ (0; 1); *τ*_*s*_[*s*] ∈ (1; 12 ∗ 3600) splines’ coefficients ∈ (−∞; ∞) sine scale [s] ∈ (1; 0.5 ∗ signal duration).

The optimization procedures were always run from multiple starting points arbitrarily set to 5 (more starting points gave no advantage) scattered uniformly over the parameter space. For simplex algorithm applied to the learning part the results typically converged to the same point implying uni-modality of the loss landscape and improving trustworthiness of the optimization procedure. We also performed some grid searches over the parameter spaces which generally agreed with the results of simplex procedure. Fitting the stochastic intensity part the obtained results were acceptable as verified by visual inspection of the timeseries or by the metrics like the Kolmogorov-Smirnov plots ^52^, even though the procedures did not necessarily converge, in the sense of reaching a given threshold of accuracy within a given number of iterations. Thorough modeling of the stochastic intensity part with proper checks using appropriate metrics, such as time-rescaling theorem and KS plots, residual analysis, and others, will be reported in the future.

Likelihoods of learning models were compared using Akaike and Bayesian Information Criteria ^30^ imposing different penalties on the number of model parameters. In view of our simulation results below we can say that the Bayesian Information Criterion better indicated the model that should be selected.

Social models were based on respective models of individual learning with additional parameters representing weight (*γ*) and time scale (*τ*_*s*_) of social influence. We considered two variants of the decay of the social influence over time, assuming exponential decay or flat memory buffer. In the first one we assumed the influence of observations decays exponentially with time, with the *τ*_*s*_ parameter being the time constant. In the second, we assumed rectangular memory buffer of the social influence with a sharp cut-off before *τ*_*s*_. Here we report results only with the exp-decay model as the results for the rectagular window did not differ significantly.

### Methods validation

To ensure that the proposed modeling and estimation framework is self-consistent and sufficiently selective to distinguish between socially interacting or non-interacting animals we simulated data with two different models and analyzed simulated data using the proposed framework. Both simulated models assumed the basic Q-learning, however one assumed the mice were agnostic of each other’s existence, the other assumed social interaction as defined above. To make the simulated data comparable with the experimental data we took the stochastic intensity from the experimental fits and simulated two populations of 14 independent or 14 interacting mice.

In the first simulation we used independent mice learning reward with the basic Q-learning with parameters *α* = 0.2, *β* = 0.85. Having simulated the trajectories we used our estimation framework to fit four non-interacting and four interacting models (the same set as for experimental data except “four alphas”). The BIC values for different model fits to individual “simulated mice” are shown in Supplementary Table 3. Fig. 7**a** shows normalized Bayesian Information Criteria (Eq. 1) for every model fit and every mouse.

**Figure 7:**
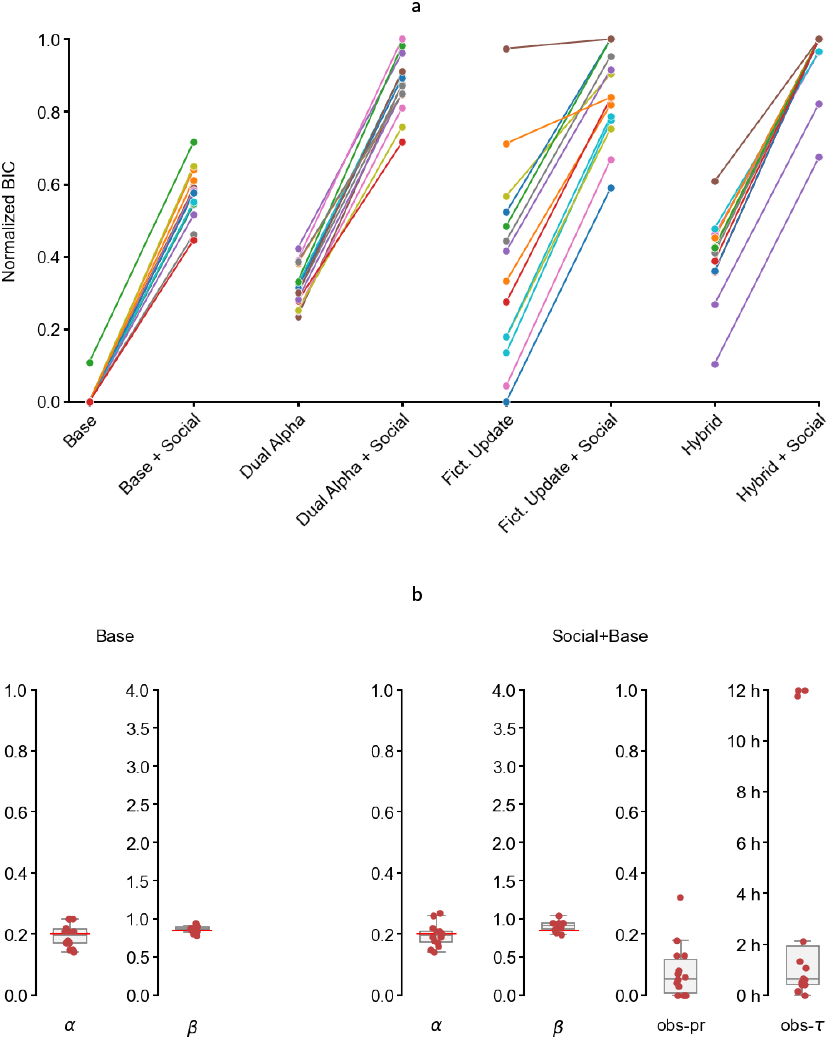
**a** Normalized Bayesian Information Criteria for every model fit and every mouse. **b** Distribution of parameter values for the base and social+base models fitted to the behavior of all the mice.

Fig. 7**b** shows fits of model parameters to behavior of mice simulated as independent. We show only fits of independent and interacting Q-learning models. Red horizontal lines indicate values assumed in the simulations *α* = 0.2, *β* = 0.85, *τ*_*s*_ = 0, *γ* = 0. We can see that the median of the distribution of *α, β* is close to the simulated values and the distribution of values for *τ*_*s*_, *γ* are not statistically significantly different from 0.

In the second simulation we assumed interacting mice learning reward with the basic Q-learning with parameters *α* = 0.35, *β* = 1 with the social component governed by *γ* = 0.55, *τ*_*s*_ = 2.2*h*. As above, having simulated the trajectories we used our estimation framework to fit the same four non-interacting and four interacting models. The BIC values for different model fits to individual “simulated mice” are shown in Table 4. Fig. 8**a** shows normalized Bayesian Information Criteria for every model fit and every mouse.

**Table 4:** Bayesian Information Criteria for model fits to simulated data. Rows: mouse number. Column: model. Highlighted are the minima for a given mouse indicating the model selected by the BIC.

| \Model<br>mouse \ | Base | Dual<br>Alpha | Fict.<br>Update | Hybrid | Base<br>+ Social | Dual Alpha<br>+ Social | Fict. Update<br>+ Social | Hybrid<br>+ Social |
| --- | --- | --- | --- | --- | --- | --- | --- | --- |
| 1 | 3741.10 | 3747.88 | 3765.65 | 3745.09 | 3700.95 | 3706.28 | 3708.06 | 3707.09 |
| 2 | 3876.97 | 3878.19 | 3892.61 | 3885.40 | 3857.25 | 3864.12 | 3864.11 | 3866.34 |
| 3 | 3828.52 | 3832.73 | 3857.65 | 3837.13 | 3792.49 | 3799.04 | 3799.31 | 3801.03 |
| 4 | 3900.40 | 3907.25 | 3915.55 | 3911.03 | 3871.10 | 3878.10 | 3874.24 | 3879.67 |
| 5 | 3730.04 | 3734.69 | 3761.62 | 3741.87 | 3683.95 | 3689.65 | 3691.22 | 3694.20 |
| 6 | 3847.64 | 3854.36 | 3864.14 | 3859.44 | 3831.56 | 3837.54 | 3836.85 | 3841.83 |
| 7 | 3779.67 | 3786.20 | 3787.65 | 3773.56 | 3763.49 | 3770.73 | 3759.79 | 3758.54 |
| 8 | 3849.73 | 3856.27 | 3877.04 | 3860.16 | 3808.34 | 3815.28 | 3815.16 | 3816.80 |
| 9 | 3789.46 | 3795.71 | 3809.77 | 3797.47 | 3759.11 | 3766.07 | 3761.38 | 3766.88 |
| 10 | 3656.68 | 3663.87 | 3669.77 | 3667.42 | 3649.33 | 3656.38 | 3653.80 | 3658.56 |
| 11 | 3780.68 | 3782.38 | 3791.20 | 3785.32 | 3756.51 | 3758.10 | 3757.75 | 3760.23 |
| 12 | 3797.18 | 3801.41 | 3803.48 | 3803.90 | 3762.53 | 3768.09 | 3762.04 | 3769.05 |
| 13 | 3697.33 | 3703.31 | 3718.64 | 3709.78 | 3667.09 | 3673.84 | 3668.70 | 3675.29 |
| 14 | 3695.05 | 3699.07 | 3696.26 | 3698.81 | 3669.03 | 3674.12 | 3665.81 | 3667.27 |
| Sum | 52970.45 | 53043.29 | 53211.03 | 53076.36 | 52572.71 | 52657.36 | 52618.23 | 52662.78 |

**Figure 8:**
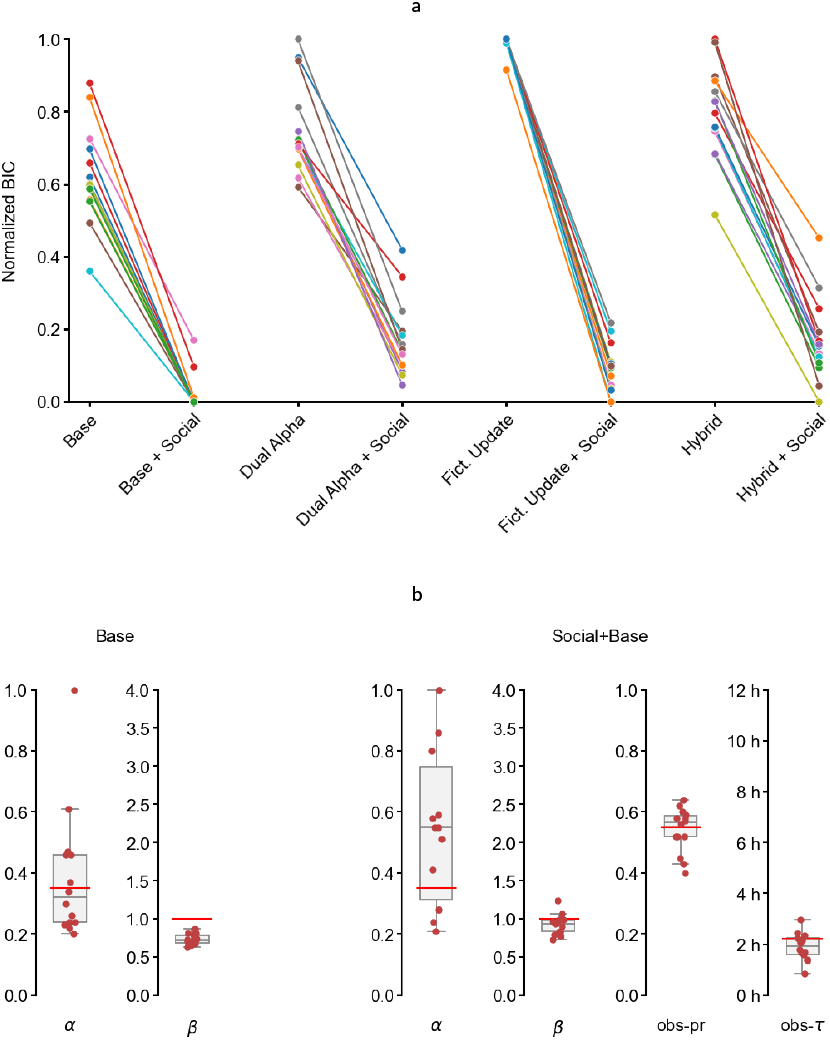
**a** Normalized Bayesian Information Criteria for every models fit and every mouse. **b** Distribution of parameter values for the base and social+base model fitted to the behavior of all the mice.

Fig. 8**b** shows fits of model parameters to the behavior of mice simulated with the social component. We show only fits of independent and interacting base Q-learning models. Red horizontal lines indicate values assumed in the simulations *α* = 0.35, *β* = 1, *τ*_*s*_ = 2.2*h, γ* = 0.55. We can see that in both fits the ground truth *α* is consistent with the distribution. When assuming independent mice, distribution of *β* is below the ground truth, which would be interpreted in experiment as more exploratory behavior but here it is a consequence of not including the following of others. The fits of the social parameters are consistent with ground truth values and clearly separated from 0.

These simulations and analysis indicate that the proposed technique is self-consistent and at least for cohorts behaving accordingly to our assumed models, with the available data size, it is sufficient to disambiguate independent and interacting mice. Selecting specific learning model is more diffficult but prevalence in a population seems robust. This is why we believe the experimentally observed high social interaction probability in previous section can be trusted. Also, the inadequacy of the simplest learning models is also reliable.

## Supplementary Materials

### BIC values for all the models for all the mice

Table 2 shows the values of the Bayesian (Schwarz) Information Criterion for the models tested ^30^. This is the negative log-likelihood that the observed data were generated by the given model (*L*) penalized by the number of parameters *k* and data points *n*: *BIC* = *k* ln *n* − 2 ln(*L*). The first column shows the animal number. The other columns present BIC values for different learning models. Interestingly, we can see that for every mouse the best model always includes social aspects. The basic Q-learning, even with social effects, is never the best explainer of the obtained data. The model which explains the data best overall is Hybrid+Social and we present the distribution of its parameters in Fig. 4. This was the best model for 6 animals but also summary BIC is the smallest among all the models. The second best is the model with four alphas which disambiguates true learning from inverse fictitious learning at the cost of extra two parameters which occasionally leads to stronger penalization but still for 4 animals this is the best model. Nevertheless, we focus our discussion of fitted models here on the Hybrid + Social model.

### BIC values for all the models fitted to simulated data from independent mice

### BIC values for all the models for all the mice fitted to simulated data from mice interacting socially

